# Circulating extracellular vesicles display immunotherapeutic targets and complement circulating tumor DNA in high-risk neuroblastoma

**DOI:** 10.64898/2026.10.21.760707

**Authors:** Martin Auber, Angelique Nejaime, Hyun-Dong Chang, René Maier, Roza Mizrak, Daniel Bachurski, Michael Hallek, Jasmin Vahid Yousefnia, Lena Andersch, Peter Kühnen, Marco Lodrini, Marvin Jens, Elke Pogge von Strandmann, Annette Künkele, Christian Preußer, Hedwig E. Deubzer

**Affiliations:** Department of Pediatric Oncology and Hematology, Charité – Universitätsmedizin Berlin, Berlin, Germany; Deutsches Rheuma-Forschungszentrum Berlin, Berlin, Germany; Department of Biochemistry, Charité – Universitätsmedizin Berlin, Berlin, Germany; Department I of Internal Medicine, Center for Integrated Oncology Aachen Bonn Cologne Duesseldorf, CECAD Center of Excellence on Cellular Stress Responses in Aging-Associated Diseases, Faculty of Medicine and University Hospital Cologne, University of Cologne, Cologne, Germany; Department of Pediatric Endocrinology and Diabetology, Charité – Universitätsmedizin Berlin, corporate member of Freie Universität Berlin and Humboldt-Universität Berlin, Berlin, Germany; German Cancer Consortium (DKTK), Partner Site Berlin, and German Cancer Research Center (DKFZ), Heidelberg, Germany‘; EV-iTEC Core Facility, ICenter for Tumor Biology and Immunology, Marburg University, Marburg, Germany; Berlin Institute of Health (BIH) at Charité, Berlin, Germany; Berlin Center for Advanced Therapies, Charité – Universitätsmedizin Berlin, Berlin, Germany; German Center for Child and Adolescent Health (DZKJ), partner site Berlin, Germany

**Keywords:** multimodal biomarker profiling, gangliosides, GD2, L1CAM, NCAM1, circulating cell-free tumor DNA, pediatric cancer

## Abstract

Extracellular vesicles (EVs) offer a minimally invasive window into tumor biology through surface molecules reflecting their cellular origin. Here, we investigated whether neuroblastoma-derived EVs retain tumor-associated surface markers and provide complementary information to established biomarkers in children with neuroblastoma. EVs from neuroblastoma cell lines were characterized using orthogonal biochemical, morphological and biophysical approaches, followed by analysis of GD2, L1CAM and NCAM1 surface expression. EV-associated markers were subsequently evaluated in blood plasma-derived EVs from children with neuroblastoma and pediatric controls and compared with clinical and circulating cell-free DNA (cfDNA) parameters. Neuroblastoma EVs displayed GD2, L1CAM and NCAM1 in patterns reflecting donor-cell surface expression, supported by a positive correlation between cellular and EV marker abundance (Spearman ρ=0.74, P<0.0001). Genetic disruption of GD2 biosynthesis resulted in corresponding loss of EV-associated GD2, further supporting its cellular origin. In plasma, GD2-, L1CAM- and NCAM1-positive EVs were enriched in patients with high-risk neuroblastoma and correlated with LDH level, soft tissue disease burden (GD2, L1CAM) and cfDNA concentration (GD2, NCAM1). Longitudinal analyses demonstrated dynamic and partially discordant changes in EV- and cfDNA-markers. These findings identify tumor-associated EV surface phenotypes as complementary circulating biomarkers and support their further evaluation for minimally invasive monitoring of neuroblastoma evolution.

## INTRODUCTION

Extracellular vesicles (EVs) are membrane-enclosed nanoparticles that mediate intercellular communication by transferring proteins, lipids, glycans and nucleic acids between cells. Derived from the plasma membrane or endosomal compartments, EVs retain aspects of the molecular composition of their cells of origin while undergoing selective cargo sorting during biogenesis.^1^ Their surface composition can capture features of the parental tumor and provide a molecular fingerprint that may be accessible through minimally invasive liquid biopsy.

In solid tumors, EVs provide a potential link between the tumor cell and the circulation. EV surface molecules can actively shape tumor progression by influencing organ tropism, extracellular matrix interactions and immune responses.^2^ Surface proteins including adhesion molecules, integrins, immune checkpoint ligands and tumor-associated antigens regulate EV uptake by recipient cells and can promote metastatic niche formation, immune suppression and therapeutic resistance. The surface of tumor-derived EVs should therefore be viewed not simply as a passive reflection of the parental cell, but as a functional interface capable of influencing the tumor microenvironment and disease progression.

Patients with the solid embryonic tumor, neuroblastoma, remain a major therapeutic challenge in pediatric oncology when stratified to be at high risk for relapse. Neuroblastoma exhibits marked clinical heterogeneity, ranging from spontaneously regressing tumors to highly aggressive and metastatic forms, which is reflected in divergent survival rates between 90– 95% for low- and intermediate-risk patients compared with only ∼50% for those with high-risk neuroblastoma.^3^ Approximately half of pediatric patients diagnosed with neuroblastoma are at high risk for relapse, with <20% survival.^4^ The incorporation of anti-disialoganglioside (GD2) immunotherapy into multimodal treatment has improved survival of high-risk neuroblastoma, establishing tumor-cell surface antigens as clinically relevant therapeutic targets.^5,6^ More recently, GD2-directed cellular therapies, including chimeric antigen receptor (CAR) T cells, have demonstrated clinical activity in relapsed or refractory high-risk neuroblastoma,^7–9^ further emphasizing the importance of tumor antigen expression and its accessibility to immune effector cells. These therapeutic advances highlight the need for minimally invasive approaches to interrogate tumor antigen biology over the course of disease.

In this study, we investigated the relationship between neuroblastoma cells and the EVs they release, focusing on the tumor-associated surface markers GD2, L1CAM and NCAM1. We asked whether the EV surface phenotype recapitulates that of the parental neuroblastoma cells and whether GD2-positive EVs can be detected in the plasma of patients with neuroblastoma. We further examined the relationship between circulating GD2-positive EVs and established clinical parameters as well as emerging cell-free DNA (cfDNA) markers. By integrating cellular, EV and circulating molecular parameters, we sought to determine whether GD2-positive EVs provide a minimally invasive readout of tumor-associated biology and whether they complement existing biomarkers for disease evolution.

## MATERIALS AND METHODS

### Patient samples

Peripheral blood samples from patients with neuroblastoma were collected as described^10^ (ethics approvals: EA2/055/17, EA2/011/23). Patients were treated at the Charité – Universitätsmedizin Berlin (high-risk, n=10; low-/intermediate-risk, n=8) and enrolled in the GPOH NB2016 Registry (n=18). Informed written patient/parent consent was obtained during registry participation. The comparative control group consisted of blood plasma from healthy children serving as hematopoietic stem cell donors for their siblings (n=9; ethics approval EA2/011/23) with a median age of 88.2 months [min-max: 35.3 – 173.3 months] and from 12 pediatric patients with compensated hypothyroidism and a median age of 81.0 months [min-max: 25.4– 151.0 months (ethics approval: EA2/131/11). Peripheral blood was centrifuged at 1,900 x g for 7 min at room temperature to separate plasma from cells.^11^ Plasma samples were centrifuged a second time at 3,250 x g for 10 min at room temperature to remove cellular debris before storage at −80 °C.^11^

### Cell culture

The human neuroblastoma cell lines, CLB-Ga (RRID: CVCL_9529), GI-ME-N (RRID: CVCL_1232), IMR-5/75 (RRID: CVCL_M473), SH-EP (RRID: CVCL_0524), SK-N-AS (RRID: CVCL_1700), SK-N-BE(2) (RRID:CVCL_0528) and SH-SY5Y (RRID: CVCL_0019) were cultured in DMEM medium (Thermo Fisher Scientific) supplemented with 10% fetal calf serum (GE Healthcare) and maintained at 37°C and 5% CO_2_. The GD2-deficient SK-N-BE(2) cell line was generated using CRISPR/Cas9 guide constructs targeting the *B4GALNT1* gene, which encodes the GD2 synthase. Guides were designed using the crispor software.^12^ Target sequences of sgRNA were: 5’-caccgCGTCCCGGGTGCTCGCGTAC-3’ (forward); 5’-aaacGTACGCGAGCACCCGGGACGc-3’ (reverse). Sequences were cloned into the CRIPSRcas9 vector pSpCas9(BB)-2A-Puro (PX459) V2.0 (Addgene). SK-N-BE(2) cells were transfected with 1 ug DNA using the Effectene transfection reagent according to the manufacturer’s instructions (Qiagen). SK-N-BE(2) cells were selected for GD2 negativity through 1 µg/ml puromycin selection followed by enrichment via flow cytometry using a BD FACSAria (BD Biosciences). Cell lines were authenticated by high-throughput SNP-based assays^13^ and regularly monitored for *Acholeplasma laidlawii*, mycoplasma species and squirrel monkey retrovirus infections using high-throughput multiplexed testing^14^. For EV isolation from conditioned medium, cells were grown to 80% confluency, then washed twice for 5 min with DMEM without supplements. Thereafter, cells were cultured for 24h in serum-free DMEM medium supplemented with 2% B27 (50x, ThermoFisher Scientific Inc.), 1% non-essential amino acids (ThermoFisher Scientific Inc.), 50 ng/ml recombinant human epithelial growth factor (Sigma-Aldrich) and 20 ng/ml recombinant human fibroblast growth factors (PeproTech Inc.). In total, 75 ml of conditioned medium per experiment were centrifuged at 200 x g for 10 min at 4 °C, then passed through a 0.45 µM cellulose acetate filter (Corning) to remove cells and larger particles.^15^

### Extracellular vesicle isolation from human blood plasma

EVs were isolated from blood plasma using the CD81 EV Isolation Kit, human (Miltenyi Biotec), which acts via positive selection of vesicles harboring the CD81 tetraspanin protein on their surfaces. In brief, 400 µL of plasma was mixed with 1600 µL of Dulbecco’s phosphate-buffered saline (Thermo Fisher Scientific Inc.).^16^ Antibody-coupled magnetic MicroBeads specifically binding CD81 were added to the mixture and incubated for 1h at room temperature. Labeled vesicle-bead samples were loaded onto μ Columns placed in the magnetic field of a µMACS^®^ Separator (Miltenyi Biotec). Magnetically labelled CD81⁺ EVs were subsequently eluted from μ Columns with 100 µL isolation buffer to a final volume of 120 µL for immediate flow cytometric analysis.

### Extracellular vesicle isolation from conditioned cell medium

Conditioned culture medium was concentrated at 4 °C approximately 40-fold using a 100K Centricon Plus-70 centrifugal filter device (Merck Millipore).^15^ Size exclusion chromatography (SEC) was performed using self-made columns with minor modifications.^17^ In brief, sepharose CL-2B (Sigma-Aldrich) columns were prepared in 10 mL syringes with a 0.22 μm polyethersulfone filter membrane placed at the syringe base, then washed with 10 mL Dulbecco’s phosphate-buffered saline (DPBS, Thermo Fisher Scientific Inc.). The concentrated conditioned medium (0.5 – 2 mL) was loaded on the column, and 1-mL fractions were subsequently collected. EV-containing fractions 4-7 were pooled and concentrated using an Amicon Ultra-2 100K centrifugal filter (Merck Millipore) to obtain a final volume of 150 - 200 µL for flow cytometry analysis.

### Western blot analysis

EV-containing samples were mixed with 4x Laemmli buffer containing 400 mM DTT, with the exception of antibodies against CD9, CD63, CD81, and calnexin (CANX). Samples were then subjected to SDS-PAGE using self-made 12% gels. Proteins were transferred to a PVDF membrane using the semidry Trans-Blot Turbo Blotting System (BioRad). Membranes were blocked with 5% milk powder and 0.1% Tween in PBS and incubated with primary antibodies followed by HRP-conjugated secondary antibodies. Chemiluminescent detection and visualization were performed to detect target proteins. If feasible, membranes were re-incubated with a primary antibody targeting a different marker protein. The following antibodies were applied: anti-CD9 (dilution 1:1000; #13403S; Abcam), anti-CD63 (dilution 1:500; #ab134045; Abcam), anti-CD81 (dilution 1:500; #MA5-13548; ThermoFisher), anti-Syntenin (SDCBP; dilution 1:10000; ab133267; Abcam), anti-Calreticulin (CALR; dilution 1:1000; #2891; Cell Signaling), anti-CANX (dilution 1:1000; #ADI-SPA-865; Cell Signaling), goat-anti-mouse-HRP (dilution 1:5000; 115–035-003; Dianova), mouse-anti-rabbit-HRP (dilution 1:5000; #5A6-1D10; Dianova).

### Transmission electron microscopy

For negative-stain transmission electron microscopy (CECAD Imaging Facility, University of Cologne), 5 µL of undiluted EV sample was adsorbed onto Formvar-coated copper grids (Science Services) for 20 min at room temperature. Grids were blocked for 30 min using Aurion blocking solution (Aurion, #905.001), then washed three times for 2 min each in PBS containing 0.1% BSA-c (pH 7.4; Aurion, #900.099). Grids were fixed with 1% glutaraldehyde in PBS for 5 min, followed by three washes in ddH₂O for 2 min each. Samples were negatively stained with 1.5% uranyl acetate (Agar Scientific, #R1260A) for 4 min before blotting and air-dried. For each sample, an unstained control grid was prepared. Images were acquired on a JEM-2100Plus transmission electron microscope (JEOL, Tokyo, Japan) operated at 200 kV, equipped with a Gatan OneView 4K camera (Gatan, Pleasanton, CA). EV size was measured using ImageJ (NIH).

### Nano-flow cytometry

EV particle concentration, size distribution and single-particle detection were analyzed using a NanoAnalyzer equipped with 488 nm and 637 nm lasers (NanoFCM Co., Ltd., Nottingham, UK). Instrument calibration, sample preparation, acquisition settings, background subtraction, and data analysis were performed as described.^18^ EVs were analyzed using a CytoFLEX nano flow cytometer (Beckman Coulter Life Sciences; 405, 488, 561, and 638 nm lasers) with CytExpert nano software. Instrument performance was verified by Daily QC and Sensitivity Monitoring using CytoFLEX nano QC Scatterspheres and Fluorospheres (Lot #5067801 and #5069801). Acquisition gains were VSSC1 60, VSSC2 189, B531 959, Y595 762, and R670 584. Events were triggered on VSSC1-H at a threshold of 366. In total,5 × 10^8^ EVs were stained in 0.1-µm-filtered PBS with CD63-AF488/GD2-APC, CD81-FITC/GD2-APC, CD81-APC/L1CAM-PE, or CD81-APC/NCAM-PE [CD63 clone H5H6; CD81 clone TAPA-1 Biolegend, GD2 NCAM L1CAM]. Samples were diluted 1:400 without washing and acquired for 30 s at 1 µL/min. Water, buffer, unstained EVs, and fluorochrome-matched isotype controls were acquired using identical settings. Background events were excluded by VSSC1-H gating followed by singlet discrimination using VSSC1-H versus VSSC1-Width. Marker-positive events were gated on the respective fluorescence channel versus VSSC1-H based on unstained and isotype controls, and identical gates were applied to corresponding samples.

### Flow cytometry of neuroblastoma cells

The immunophenotyping was conducted on unfixed neuroblastoma cells as described^19^ using the following fluorophore-conjugated antibodies: mouse anti-human GD2 monoclonal antibody (clone 14.G2a; BD Biosciences) conjugated to Brilliant Violet 421 (BV421); the recombinant human L1CAM monoclonal antibody (clone REA163; Miltenyi Biotec) conjugated to allophycocyanin (APC); and the mouse anti-human CD56 monoclonal antibody (clone HCD56; BioLegend) conjugated to phycoerythrin (PE). Dead cells were excluded using the LIVE/DEAD® Fixable Near IR Dead Cell Stain Kit (ThermoFisher Scientific Inc.). Flow cytometry was conducted on a LSRFortessa X-20 (BD), and data were analyzed using the FlowJo v10 software (Tree Star Inc). The signal obtained from unstained cells was used to measure the background autofluorescence and served as the baseline for the stained samples to yield background-corrected cell-surface antigen fluorescence data relative to this baseline.

### Detecting extracellular vesicle surface epitopes

Surface proteins/glycolipids were detected on extracellular vesicles using the MACSPlex EV Kit Neuro, human (Miltenyi Biotec), which allows detection of 37 surface epitopes and 2 isotype controls according to the manufacturer’s instructions. Briefly, 120 µL of sample containing the purified extracellular vesicles (from blood plasma or conditioned cell culture medium) or buffer control (DPBS) were incubated overnight with the capture beads. Following a washing step, the samples were stained with a combination of detection antibodies (CD9-, CD63- and CD81-APC). After further washing steps, flow cytometry analysis was performed using the MACSQuant^®^ Analyzer 16 (Miltenyi Biotec) and the MACSQuantify^®^ software (v2.13.3). The 39 bead populations were distinguished through different fluorescence intensities detected by the fluorescein isothiocyanate (FITC, B1) and PE (B2) channels. The signal intensity of each epitope in the sample was analyzed as fold change of the respective signal intensity measured in the buffer control sample.^20^ To minimize operator bias, a signal intensity threshold was established for each of the 3 tetraspanins (CD9, CD63, CD81) employed as well-established markers for extracellular vesicles.^21–23^ We used the previously established method^23^ for semi-quantitative GD2, L1CAM and NCAM1 analysis, where at least 2 of the 3 tetraspanins analyzed must have had a signal intensity >2 to include the sample for further analysis.

### Statistical analysis

Levels of GD2-, L1CAM- and NCAM1-positive extracellular vesicles in blood plasma samples were first normalized to controls (log10 ratio of measured level to buffer) and then compared among the different patient groups using the non-parametric two-tailed Mann-Whitney U test to evaluate significance. P-values were adjusted to control for multiple hypothesis testing using the Benjamini-Hochberg method, and the resulting adjusted values are reported as Q-values.^24^ The statistical relationship between two variables was calculated using Spearman rank correlation as indicated in the text. All tests were conducted using GraphPad Prism^®^ version 10.4.2.

### EV-TRACK

All relevant experimental data have been submitted to the EV-TRACK knowledgebase^25^ (EV-TRACK ID: EV250106).

## RESULTS

### Neuroblastoma sEVs display GD2, L1CAM and NCAM1 in proportion to donor-cell surface expression

Conditioned medium from neuroblastoma cell cultures was processed by SEC for phenotypic analyses in the *in vitro* experimental study (**Suppl. Fig. S1A**). The conditioned medium was collected, cleared by centrifugation and filtration, concentrated and subjected to SEC to obtain the EV-containing fractions 4-7. The quality of the isolated EV preparations was assessed using complementary phenotypical, morphological and biophysical approaches as demonstrated for EVs derived from IMR-5/75 cells (**Fig. 1**). Western blotting demonstrated enrichment of the canonical EV-associated proteins CD9, CD63, CD81 and the SDCBP binding protein, whereas the endoplasmic-reticulum proteins CANX and CALR were not detected in the EV fraction, supporting the absence of substantial cellular contamination (**Fig. 1A**). Transmission electron microscopy independently confirmed the presence of membrane-enclosed vesicles with the characteristic cup-shaped appearance (**Fig. 1B**). Nano-flow cytometry showed a particle population with a mean size peak of 73.5 nm, consistent with the expected size range of small EVs (sEVs; **Fig. 1C**). In accordance with the Minimal Information for Studies on Extracellular Vesicles (MISEV) guidelines^26^, these orthogonal analyses together established that the SEC-based workflow reproducibly yielded preparations enriched in sEVs suitable for downstream surface-marker analyses.

**Fig. 1.**
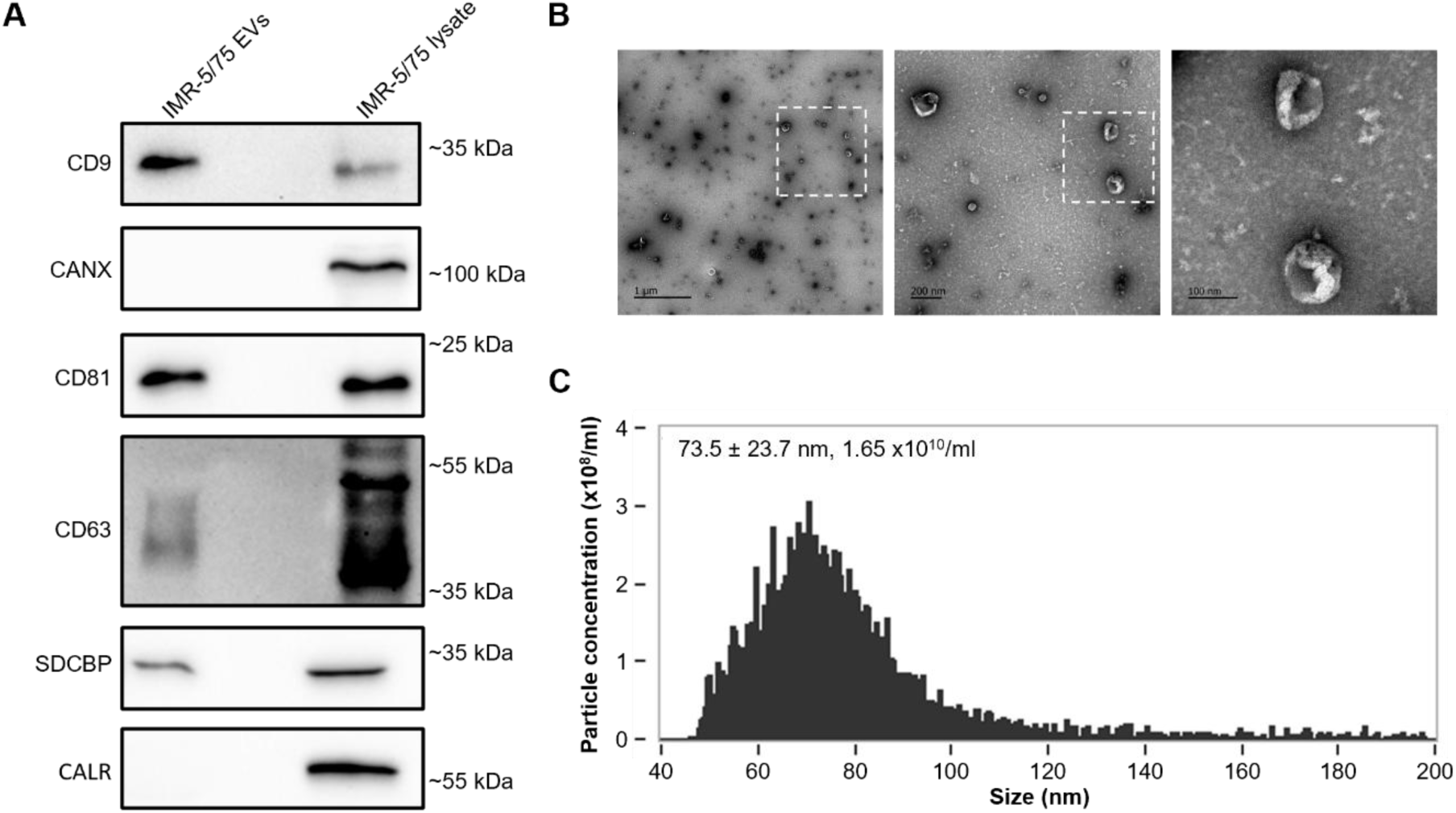
Quality control of size-exclusion chromatography-isolated small extracellular vesicles (sEVs) used for *in vitro* experiments. sEV preparations obtained from IMR-5/75 neuroblastoma cells were assessed using three complementary approaches: **A**, western blotting for EV and cellular markers for purity; **B**, transmission electron microscopy for EV size and morphology; and **C**, nano-flow cytometry for EV concentration and size distribution.

We next investigated whether neuroblastoma-derived sEVs carry surface molecules that reflect the phenotype of their parental cells. sEVs isolated from six neuroblastoma cell lines (**Fig. 2A**) were analyzed using the MACSPlex-based flow cytometry.^23^ The EV markers CD63 and CD81 were consistently detected across all sEV preparations, while CD9 was detectable in sEVs from most cell lines, albeit at lower levels in GI-ME-N derived sEVs (**Fig. 2B**). Phenotypic differences observed between sEV preparations were thus assessed in the context of sEV populations that were uniformly positive for established EV-associated markers. Marked differences were observed in the abundance of GD2, L1CAM and NCAM1 on sEVs derived from the different neuroblastoma cell lines. GD2 was particularly prominent on sEVs from SH-EP, SH-SY5Y, IMR-5/75 and CLB-Ga cells, whereas SK-N-AS-derived sEVs showed an intermediate GD2 signal and GI-ME-N-derived sEVs only displayed a weak GD2 signal (**Fig. 2B**). These differences broadly paralleled the GD2 phenotype of the corresponding donor cells. SH-EP, SH-SY5Y, IMR-5/75, and CLB-Ga cells were strongly GD2-positive, whereas SK-N-AS and GI-ME-N cells showed only weak GD2 positivity. (**Fig. 2A**). Thus, the relative abundance of GD2 on sEVs followed the GD2 expression pattern of the cells from which the sEVs originated, ranging from low abundance in the GD2-low cell lines to strong EV-associated signals in GD2-high cell lines. A similar relationship was observed for the neuroblastoma-associated surface molecules L1CAM and NCAM1. Although their relative abundance varied between cell lines, both proteins were readily detectable on sEVs, indicating that neuroblastoma sEVs carry multiple cell surface-associated molecules rather than GD2 alone (**Fig. 2A**). The comparison of cellular and sEV phenotypes suggested that the molecular composition of the sEV surface may, at least in part, retain information about the surface phenotype of the donor cell.

**Fig. 2.**
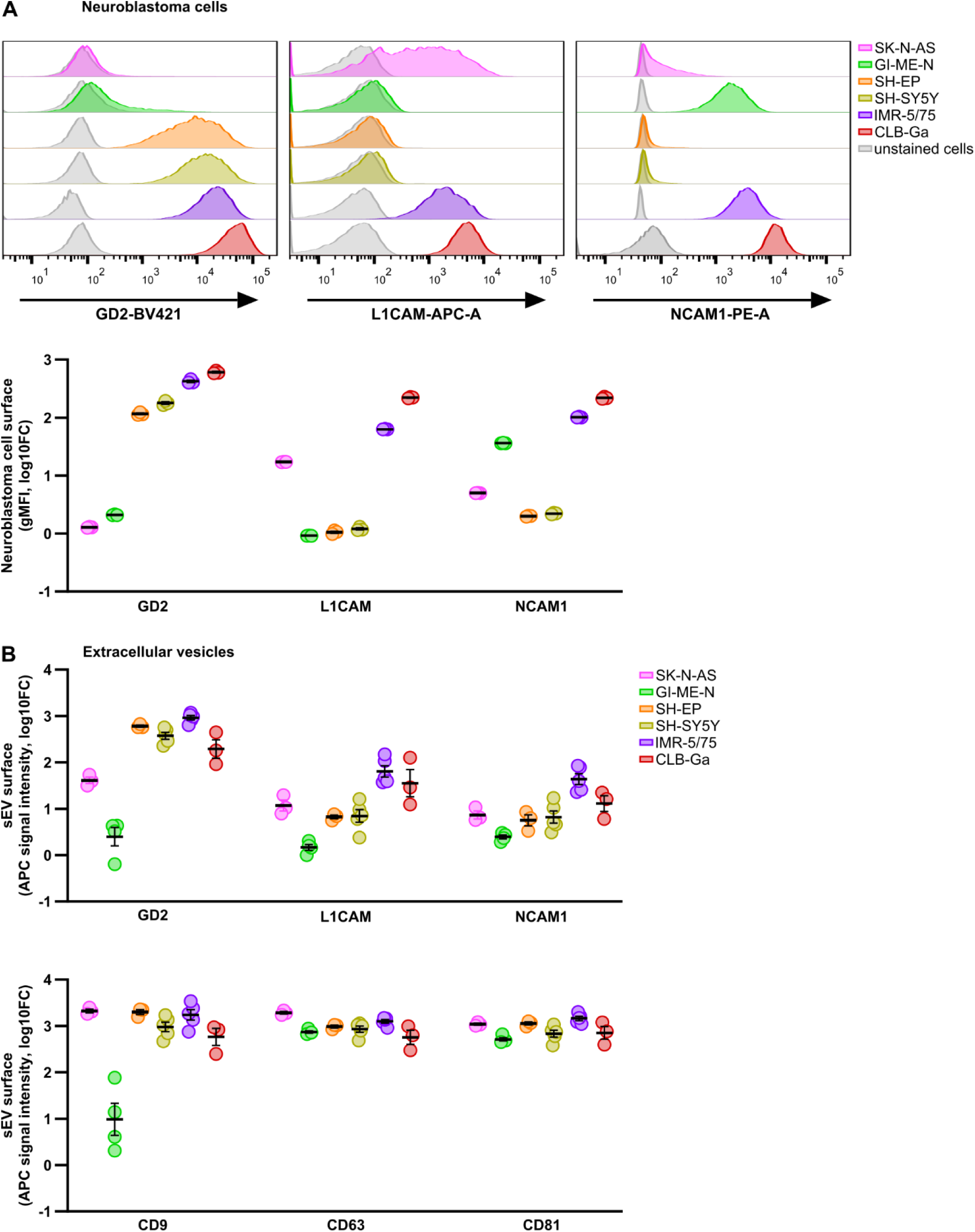
Flow-cytometric analysis of surface markers on neuroblastoma cells and small extracellular vesicles. **A**, Flow-cytometric analysis of surface GD2, L1CAM and NCAM1 expression on the indicated human neuroblastoma cell lines. Representative fluorescence histograms are shown with unstained cells as controls. Quantification is presented as geometric mean fluorescence intensity (gMFI), expressed as log10-fold change relative to the respective unstained control. **B**, Flow-cytometric analysis of GD2, L1CAM, NCAM1, CD9, CD63 and CD81 on small extracellular vesicles (sEVs) isolated from conditioned medium after 24 h of culture of the indicated neuroblastoma cell lines. Data are shown as log10-fold change in APC fluorescence intensity relative to buffer control. Individual measurements and arithmetic mean ± S.E.M. are shown.

To directly test whether sEV-associated GD2 depends on GD2 expression by the originating cell, we generated a genetically defined GD2-deficient model using CRISPR/Cas9-mediated disruption of the *B4GALNT1* gene encoding the GD2 synthase. Flow cytometric analysis confirmed loss of GD2 surface expression in *B4GALNT1* knock-out SK-N-BE(2) cells, whereas wildtype SK-N-BE(2) cells retained GD2 expression (**Fig. 3A**). Analysis of sEVs released by these cells revealed a corresponding loss of sEV-associated GD2 in the *B4GALNT1* knock-out condition, while sEVs from wildtype cells remained strongly GD2-positive (**Fig. 3B**). The canonical tetraspanins CD9, CD63 and CD81 remained robustly detected on sEVs from both genotypes, indicating that disruption of *B4GALNT1* and the resulting loss of GD2 did not substantially impairs EV release or eliminate the sEV population. The genetic experiment further helped distinguish selective changes in sEV surface composition from a general alteration in sEV production. Whereas GD2 was strongly reduced both on the *B4GALNT1* knock-out cells and on their sEVs, L1CAM and NCAM1 remained detectable on the knock-out cells and were also retained on EVs (**Fig. 3A-B**). These findings altogether demonstrate that the loss of GD2 from the EV surface specifically reflects the loss of GD2 biosynthesis in the donor cell rather than a broad disruption of EV biogenesis or release. We integrated the measurements obtained across the neuroblastoma cell lines to determine whether cell-surface expression was quantitatively associated with EV-associated marker abundance. Correlation analysis across 24 measurements demonstrated a significant positive association between marker abundance on the donor-cell surface and the corresponding EV signal (Spearman ρ = 0.73, P < 0.0001; **Fig. 3C**). The relationship was evident across the cell line panel and was further supported by the genetically defined SK-N-BE(2) wildtype and *B4GALNT1* knock-out pair. To complement the MACSPlex sEV bulk analyses, we performed single vesicle phenotyping. While L1CAM and NCAM1 were not reliably detectable at single EV level (**Suppl. Fig. S2**), both markers were detectable by bulk MACSPlex analysis (**Fig. 2B**, **3B**) as well as by western blot analysis in a previous study^27^. GD2 was consistently detectable at single sEV resolution in IMR-5/75 and SK-N-BE(2) wild type-derived populations, whereas it was not detectable in SK-N-BE(2) *B4GALNT1* knockout-derived sEV-populations (**Fig. 4A-B**). Dual staining demonstrated varying CD63^+^ and CD81^+^ positivity in the GD2^+^ sEV population derived from IMR-5/75 and SK-N-BE(2) wildtype cells, pointing towards heterogeneous surface compositions in GD2^+^ sEV-populations (**Fig. 4A-B**). Together, these findings support the concept that neuroblastoma EVs retain measurable information about the surface phenotype of their cells of origin and, in particular, identify EV-bound GD2 as a potential surrogate marker of GD2 expression in neuroblastoma cells.

**Fig. 3.**
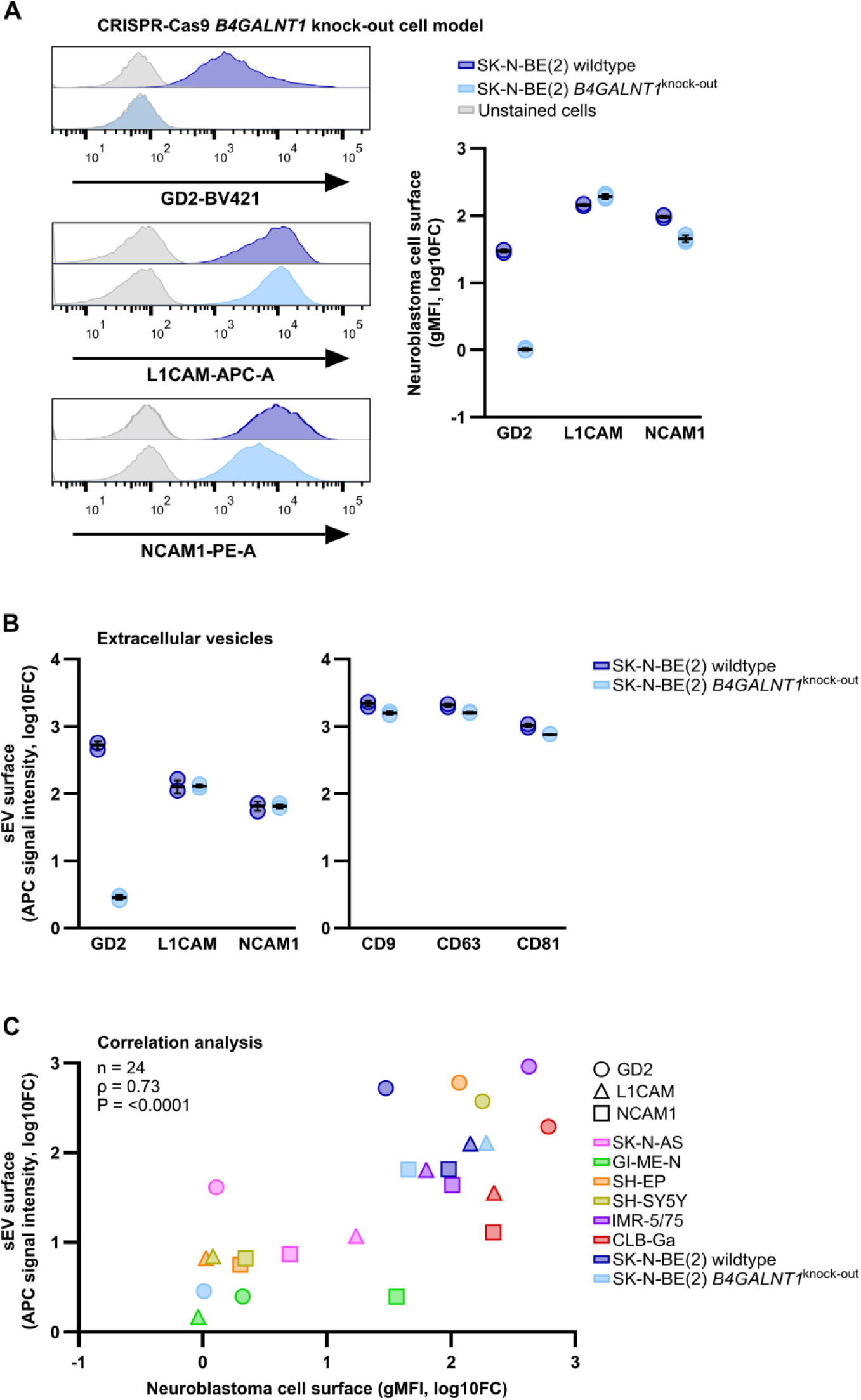
Flow-cytometric analysis of surface markers on SK-N-BE(2) cells and small extracellular vesicles in a *B4GALNT1* knock-out model. **A,** Flow-cytometric analysis of surface GD2, L1CAM and NCAM1 expression on SK-N-BE(2) wildtype and *B4GALNT1* knock-out cells. Representative fluorescence histograms are shown with unstained cells as controls. Quantification is presented as geometric mean fluorescence intensity (gMFI), expressed as log10-fold change relative to the respective unstained control. **B,** Flow-cytometric analysis of GD2, L1CAM, NCAM1, CD9, CD63 and CD81 on small extracellular vesicles (sEVs) isolated from culture medium after 24 h of culture of SK-N-BE(2) wildtype and *B4GALNT1* knock-out cells. Data are shown as log10-fold change in APC fluorescence intensity relative to buffer control. Individual measurements and arithmetic mean ± S.E.M. are shown. **C,** Correlation analysis of cell-surface marker expression and the corresponding marker expression on sEVs across the indicated neuroblastoma cell lines and SK-N-BE(2) wildtype and *B4GALNT1* knock-out cells (n = 24). Cell-surface expression is plotted as gMFI (log10-fold change) against sEV marker expression as APC signal intensity (log10-fold change). Symbols indicate GD2, L1CAM and NCAM1. Spearman correlation analysis was performed.

**Fig. 4.**
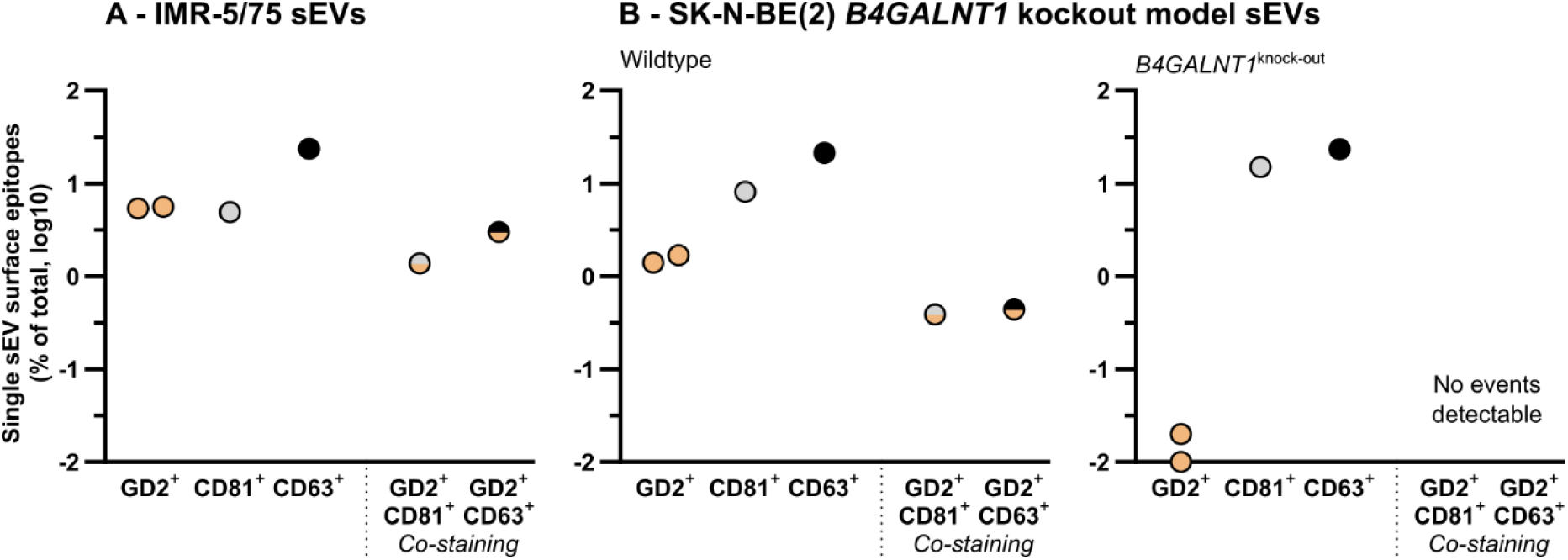
Flow-cytometric analysis of surface markers on single small extracellular vesicles derived from neuroblastoma cell models. Small EVs isolated from conditioned cell culture medium from IMR-5/75 cells (**A**) and the SK-N-BE(2) *B4GALNT1* knock-out model (**B**; *left panel*, wildtype; *right panel*, *B4GALNT1* knock-out) were subjected to dual staining using fluorophore-conjugated antibodies against GD2 (APC), CD81 (FITC) and CD63 (AF488). Representative single-event data are also shown. Fluorescence intensities obtained with different fluorophores are not directly comparable due to differences in their brightness and fluorescence properties. The percentages of positive events are shown per sample (dot plot presentation) and per condition.

### Circulating GD2-, L1CAM- and NCAM1-positive EVs associate with soft-tissue disease burden in high-risk neuroblastoma

Having established that EV-associated GD2, L1CAM and NCAM1 reflects the surface phenotype of the donor cells, we investigated whether GD2^+^-, L1CAM^+^- and/or NCAM1^+^-EVs can be detected in the circulation of children with neuroblastoma. Only pretreatment samples were included in this single time point analysis. As the complex composition of blood plasma can lead to the co-isolation of non-EV components when using size- or density-based bulk EV isolation strategies^28–30^ that are suitable for the isolation of sEVs from conditioned medium (**Suppl. Fig. S1A**), we used an immunoaffinity-based EV isolation strategy using magnetic beads (**Suppl. Fig. S1B**). A pilot experiment comparing anti-CD9, anti-CD63 or anti-CD81 coated beads in combination with different volumes of blood plasma from a patient with high-risk neuroblastoma demonstrated that anti-CD81 coated beads resulted in the comparably highest detection of the tumor-associated EV surface markers GD2, L1CAM and NCAM1 (**Suppl. Fig. S3**). Based on these findings, and given the co-detection of CD81 and GD2 on individual EV events by single vesicle analysis (**Fig. 4**), CD81-based bead isolation was selected for subsequent plasma EV analyses. As plasma volume correlated with EV surface marker detection (**Suppl. Fig. S3**), we used the highest possible volume (400 µl) for all subsequent comparative analyses. The clinical cohort comprised patients with high-risk neuroblastoma, patients with low- or intermediate-risk neuroblastoma and pediatric controls (**Fig. 5A**). Because CD81 was used for capture, its signal intensity could not be independently quantified in the clinical cohort. CD9 and CD63 remained readily detectable and served as canonical EV-associated markers (**Fig. 5B**). The control cohort included healthy children and children with compensated hypothyroidism. The subgroups displayed comparable EV marker profiles, with no detectable differences in CD9, CD63, GD2, L1CAM or NCAM1 signals (**Suppl. Fig. S4**). The combined pediatric control cohort thus provided an appropriate comparator for subsequent analyses of circulating EV phenotypes. Consistent detection of CD9 and CD63 across clinical groups confirmed successful recovery of EVs from plasma (**Fig. 5B**). The abundance of disease-associated surface molecules markedly differed between groups. EV-associated GD2 was strongly enriched in plasma from patients with high-risk neuroblastoma compared with low-/intermediate-risk neuroblastoma and pediatric controls (**Fig. 5B**). The high-risk neuroblastoma group also displayed increased EV-associated L1CAM and NCAM1 signals relative to all other groups. In summary, the circulating EV compartment in children with newly diagnosed high-risk neuroblastoma is characterized by a distinct surface phenotype enriched for neuroblastoma-associated molecules, confirming the observations made *in vitro* in the clinical setting.

**Fig. 5.**
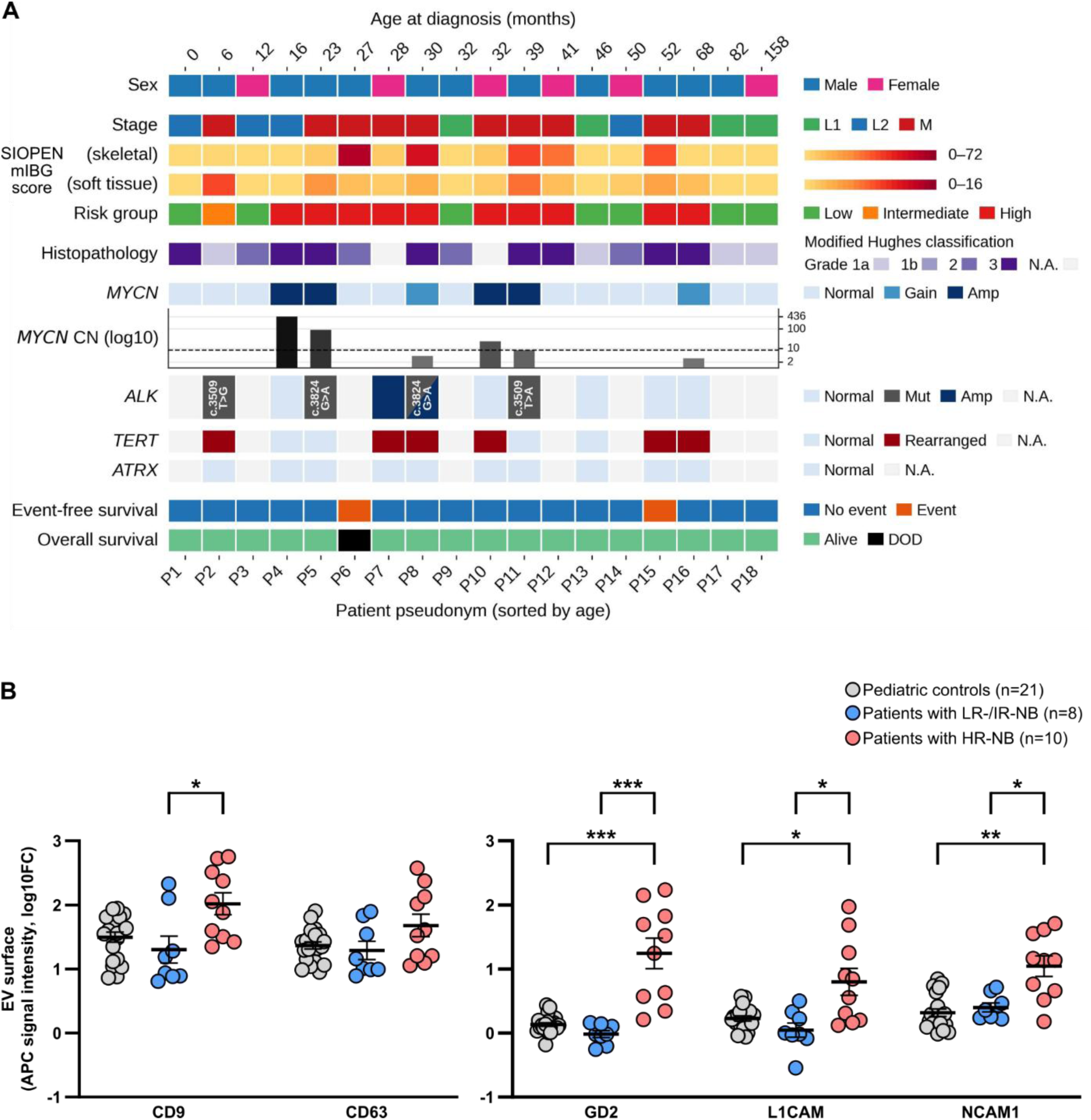
Clinical characteristics and extracellular vesicle surface-marker analysis in pediatric patients with neuroblastoma. **A**, Clinical and molecular characteristics of the neuroblastoma cohort, including age at diagnosis, sex, stage, skeletal and soft-tissue metaiodobenzylguanidine (mIBG) scores, risk group, histopathology, *MYCN* status and copy number (CN), *ALK* status, *TERT* rearrangement status, *ATRX* status, event-free survival and overall survival. Patients are shown with pseudonyms and ordered by age at diagnosis. N.A., not available; Amp, amplification; Mut, mutation; DOD, died of disease. **B**, Flow-cytometric analysis of CD9, CD63, GD2, L1CAM and NCAM1 on extracellular vesicles isolated from human blood plasma using CD81-targeting magnetic beads from pediatric controls (n = 21), patients with low- or intermediate-risk neuroblastoma (LR/IR-NB; n = 8) and patients with high-risk neuroblastoma (HR-NB; n = 10). Data are shown as log10-fold change in APC signal intensity relative to control, with individual measurements and arithmetic mean ± S.E.M. *Q < 0.05; **Q ≤ 0.01; ***Q ≤ 0.001 (two-tailed Mann-Whitney U test with Benjamini-Hochberg correction).

We next asked whether EV-associated GD2, L1CAM and NCAM1 were related to clinical and molecular indicators of neuroblastoma (**Fig. 6**). EV marker abundance showed distinct correlation patterns with biochemical markers. Neuron specific enolase (NSE) levels significantly correlated with NCAM1^+^ EVs (ρ = 0.59, Q = 0.03), whereas the association with GD2^+^ and/or L1CAM^+^ EVs did not reach statistical significance. LDH levels showed significant positive correlations with GD2 (ρ = 0.75, Q = 0.012) and NCAM1 (ρ = 0.64, Q = 0.024) whereas L1CAM did not show any correlation (**Fig. 6**). These findings indicate that increased circulating EV-associated marker abundance is associated with biochemical measures of tumor burden or disease activity, although the strength of these relationships differs between individual EV markers. Vanillylmandelic acid (VMA) secreted in urine did not show significant correlations with GD2^+^-, L1CAM^+^- or NCAM1^+^-positive EVs in plasma (**Fig. 6**). Homovanillic acid (HVA) in urine showed a significant correlation with GD2^+^ EVs (ρ = 0.61, Q = 0.024), whereas correlations with L1CAM^+^- and NCAM1^+^-EVs did not reach statistical significance. The association of EV markers with metaiodobenzylguanidine (mIBG) imaging-derived scores^31^ was similarly marker-dependent. For the skeletal mIBG SIOPEN score, all three markers were not statistically significant. In contrast, the EV-associated markers GD2 and L1CAM correlated with the soft-tissue mIBG SIOPEN score: GD2, ρ = 0.84 (Q = 0.018); and L1CAM, ρ = 0.83 (Q = 0.019) whereas NCAM1 didn’t reach statistically significant (**Fig. 6**). Together, these data suggest that the EV phenotype is associated with selected measures of disease burden, particularly LDH and soft-tissue disease involvement, rather than representing a uniform surrogate for every established neuroblastoma biomarker.

**Fig. 6.**
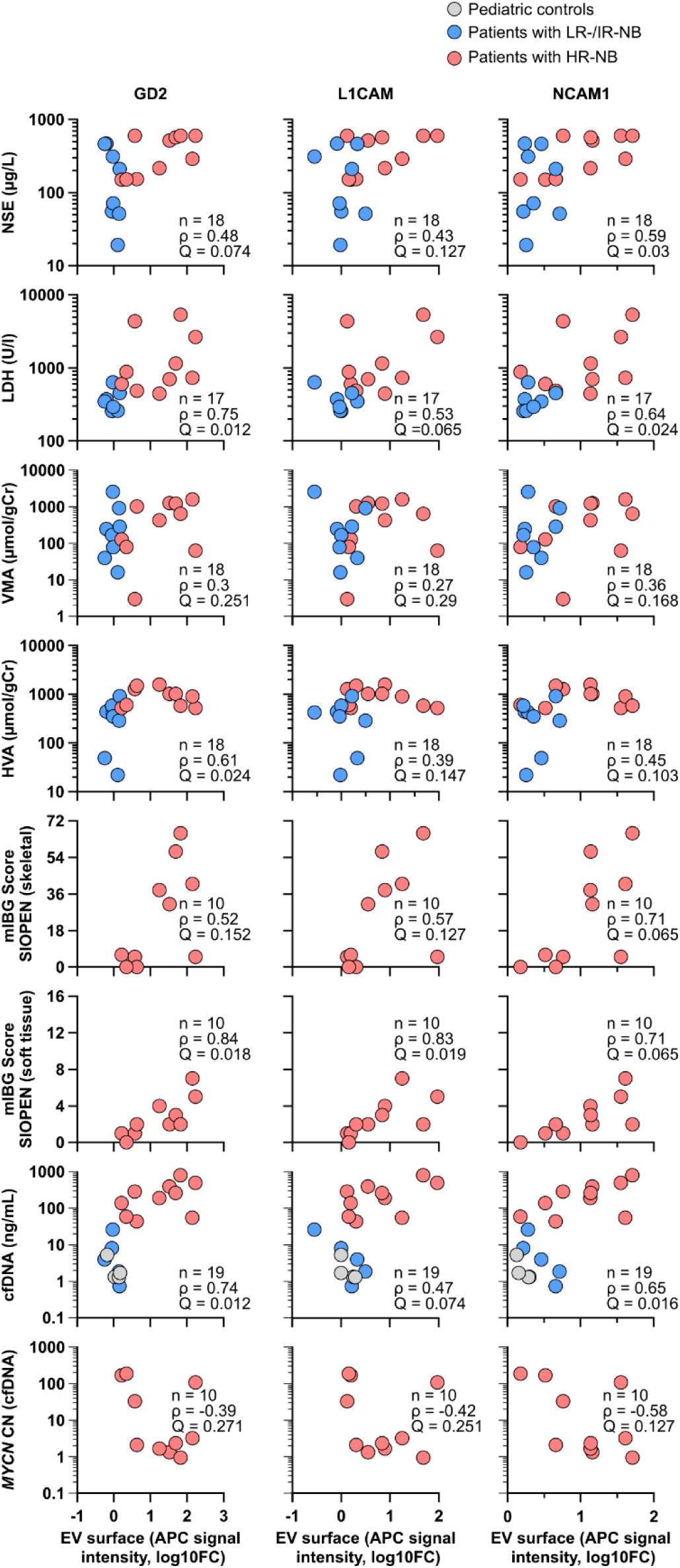
Correlation of extracellular vesicle surface-marker expression with clinical and molecular parameters in pediatric patients with neuroblastoma. Spearman correlation analysis of extracellular vesicle GD2, L1CAM and NCAM1 expression with serum neuron-specific enolase (NSE), lactate dehydrogenase (LDH; units/l), urinary vanillylmandelic acid (VMA) and homovanillic acid (HVA), metaiodobenzylguanidine (mIBG) scores for skeletal and soft-tissue involvement, circulating cell-free DNA (cfDNA) concentration and *MYCN* copy number (CN). EV marker levels are expressed as log10-fold change in APC signal intensity. Spearman correlation coefficients (ρ) and Benjamini-Hochberg corrected two-sided Q-values are shown. n indicates the number of samples analyzed for each correlation. VMA and HVA concentrations are normalized to urinary creatinine (gCr).

A notable finding was the relationship between EV surface markers GD2 and NCAM1 with circulating cfDNA. These EV-associated markers showed significant positive correlations with cfDNA concentration: GD2 (ρ = 0.74, Q = 0.012) and NCAM1 (ρ = 0.65, Q = 0.016) whereas L1CAM did not show significant correlation (**Fig. 6**). EV marker abundance did not significantly correlate with *MYCN* copy number measured in cfDNA (**Fig. 6**). The absence of a relationship with *MYCN* copy number, despite significant associations with total cfDNA concentration, indicates that EV-associated markers provide information that is not simply redundant with this specific molecular characteristic of the tumor.

### Longitudinal dynamics of EV-associated markers provide complementary information to cell-free DNA

We next examined whether changes in circulating EV markers tracked longitudinally mirrored blood-based total cfDNA dynamics in individual patients. The longitudinal analysis also incorporated molecular markers of tumor genotype, including *MYCN* and *ALK* copy number as well as the ALK p.R1275Q variant. Longitudinal measurements from three patients of the pilot cohort (**Fig. 5A**) demonstrated dynamic changes in EV-associated GD2, L1CAM and NCAM1 over time. In *Patient P5*, the EV marker profile substantially declined after the pretreatment time point TP1, with GD2, L1CAM and NCAM1 increasing during therapy at TP4 before declining again at later sampling (**Fig. 7A**). These changes occurred alongside a pronounced reduction in cfDNA concentration, normalization of the cfDNA-derived *MYCN* and *ALK* copy numbers and negativity for the ALKp.R1275Q mutation detected with 5% MAF at TP1. Similarly, in *Patient P8*, EV-associated markers remained comparatively low after the initial pretreatment sampling and showed limited variation during follow-up, whereas cfDNA decreased markedly before increasing again at the latest time point (**Fig. 7B**). In *Patient P7*, the EV markers showed relatively stable or modestly changing profiles despite a progressive decrease in cfDNA concentration and normalization of the cfDNA-derived *MYCN* and *ALK* copy numbers (**Fig. 7C**). These patient-level trajectories illustrate that EV-associated markers and cfDNA can show related but non-identical temporal behavior. The profiles of this pilot study suggest that EV-associated surface markers constitute a complementary circulating readout that can change over time in parallel with, but not necessarily concordant with conventional cfDNA-based measurements. Taken together, these findings extend the observations from neuroblastoma cell lines to the clinical setting and identify a distinct circulating EV phenotype in children with high-risk neuroblastoma. The dynamic and partially discordant trajectories observed support the potential of EV profiling as a complementary component of cfDNA analysis in patients with high-risk neuroblastoma.

**Fig. 7.**
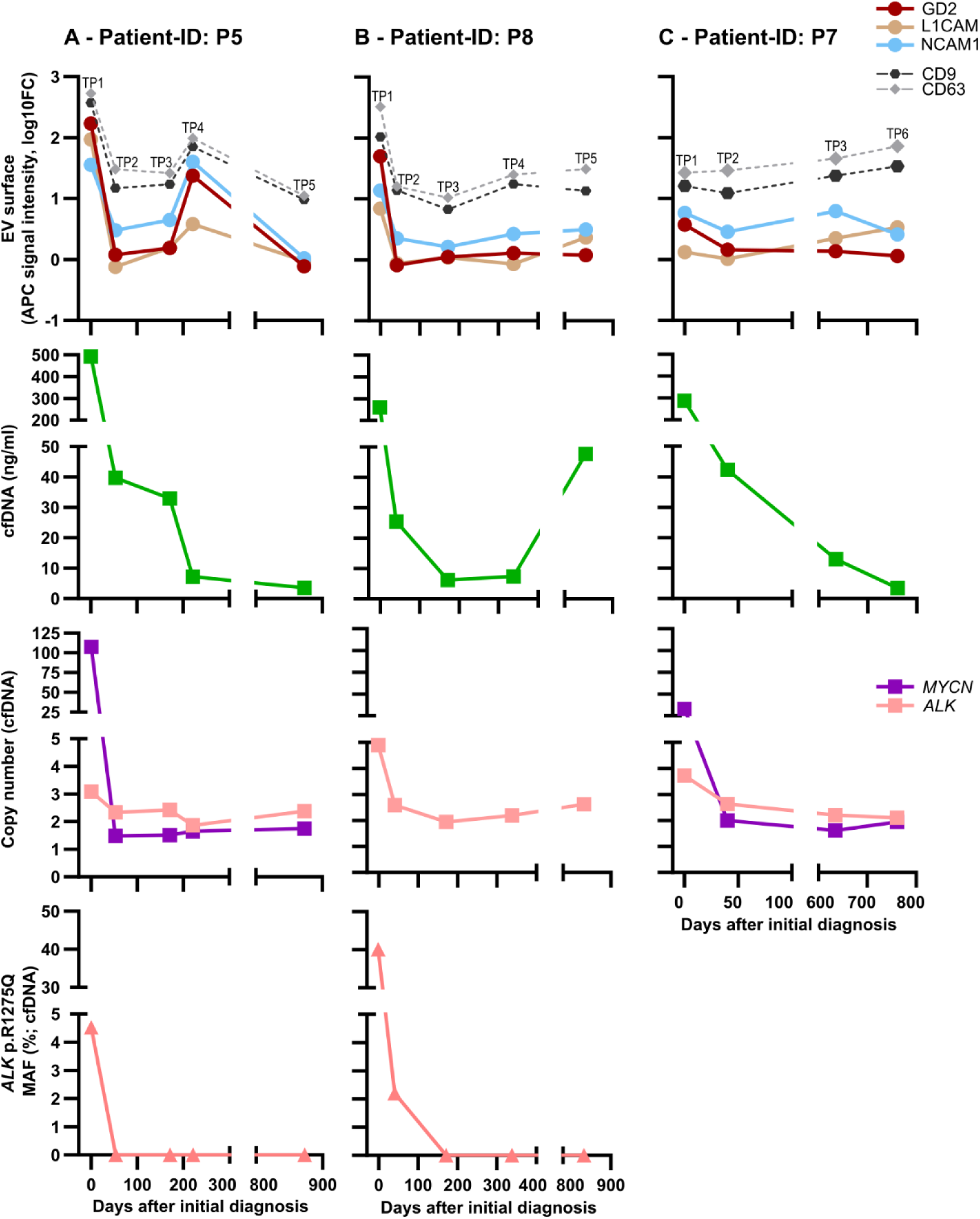
Serial assessment of extracellular vesicle and cell-free DNA parameters in patients with high-risk neuroblastoma. **A**-**C**, Longitudinal measurements in patients P5, P8 and P7, respectively, at the indicated time points after initial diagnosis. Extracellular vesicle (EV) surface-marker levels for GD2, L1CAM, NCAM1, CD9 and CD63 are shown as log10-fold change in allophycocyanin (APC) signal intensity. Cell-free DNA (cfDNA) concentration, *MYCN* and *ALK* copy number, and the mutant allele frequency (MAF) of the *ALK* p.R1275Q variant in cfDNA are shown for the corresponding time points. Values are connected between consecutive sampling time points to illustrate longitudinal changes within each patient. TP1, time point initial diagnosis, pretreatment; TP2, mid-induction chemotherapy; TP3, end of induction chemotherapy; TP4, pre-maintenance; TP5, end of treatment; TP6, follow-up.

## DISCUSSION

In this study, we demonstrate that neuroblastoma-derived EVs retain a characteristic tumor-associated surface phenotype comprising the membrane proteins L1CAM and NCAM1, as well as the clinically relevant ganglioside GD2. Neuroblastoma EV phenotyping has largely focused on generic EV markers, protein and RNA cargo, or single EV-associated tumor markers. Here, we highlight the relevance of EV membrane composition by (i) characterizing the tumor-associated surface molecules GD2, L1CAM, and NCAM1 on EVs, (ii) correlating the EV-marker abundance with the marker expression on the corresponding NB donor cells, and (iii) relating plasma EV surface profiles to clinical and molecular indicators of neuroblastoma. Our findings extend current concepts of EV phenotyping by demonstrating that tumor-associated surface molecules, including a membrane glycolipid, are preserved on neuroblastoma-derived EVs. These observations support the concept that EV membranes retain biologically informative features of their parental cells.

GD2 is particularly relevant in this context because of its distinct biochemical properties. Unlike L1CAM and NCAM1, which are transmembrane glycoproteins, GD2 is a disialoganglioside embedded within the outer leaflet of the plasma membrane. Our results demonstrate that L1CAM and NCAM1 were detectable in bulk EV analyses but not reliably at single-EV level, whereas GD2 was consistently detectable on individual EVs. This discrepancy may reflect differences in surface marker abundance, with GD2 potentially present at higher density on EVs than L1CAM and NCAM1. This difference is biologically important because EV formation involves regulated processes of membrane remodeling.^32^ EV membranes are enriched in cholesterol, sphingomyelin, ceramide and glycosphingolipids, and lipidomic studies have demonstrated that EVs exhibit a distinct lipid composition compared with parental plasma membranes, supporting budding from ordered membrane domains during vesicle biogenesis.^33,34^ Thus, the presence of GD2 on neuroblastoma-derived EVs may reflect the preservation of specialized membrane domains during EV formation.

The relationship between GD2 biology and EV biogenesis is further supported by the intracellular pathways responsible for ganglioside synthesis and trafficking. GD2 is primarily synthesized in the Golgi apparatus^35–37^ and the functional connection between Golgi-derived membrane trafficking and the endosomal system provides a plausible mechanism for GD2 incorporation into intraluminal vesicles and subsequent exosomal release.^32,38^ In addition, GD2 is associated with lipid rafts^39^, membrane microdomains involved in membrane organization and vesicle formation.^40^ Since GD2 is also present on the plasma membrane of neuroblastoma cells, incorporation into EV populations originating from the cell surface, including ectosomes, is also likely. Together, these observations support a model in which EV-associated GD2 reflects physiological membrane organization and trafficking processes.

The detection of GD2 on EVs expands the concept of EV surface profiling beyond conventional protein markers and identifies the tumor surface glycolipidome as a previously underexplored component of EV biology. Unlike soluble antigens generated through membrane turnover or proteolytic cleavage, EV-associated glycolipids remain embedded within a lipid bilayer and maintain their membrane-associated configuration. This may enable highly specific antibody-based detection approaches by preserving native antigen presentation. GD2 is particularly attractive in this context because it is expressed by the majority of neuroblastomas while showing restricted expression in normal tissues, contributing to its established role as a highly tumor-associated therapeutic target.^41,42^ Its clinical relevance is demonstrated by the successful incorporation of anti-GD2 immunotherapy into neuroblastoma treatment strategies.^42,43^

Although soluble GD2 has previously been investigated as a circulating biomarker for neuroblastoma^44,45^, our findings demonstrate that GD2 can also be detected in an EV-associated form in plasma from patients with neuroblastoma. This distinction may be biologically important because the amphiphilic nature of GD2 raises questions regarding its organization in circulation. Additional studies will be required to determine whether circulating GD2 is exclusively associated with EV membranes or whether other forms, including micellar structures or protein-associated complexes, contribute to plasma GD2 measurements. Nevertheless, these findings establish EV-associated GD2 as an additional representation of the neuroblastoma surface phenotype detectable through liquid biopsy approaches.

Using a stringent and previously validated plasma EV isolation workflow^20,23,46^, we identified enrichment of GD2-positive EVs in patients with high-risk neuroblastoma compared with healthy controls and patients with low- or intermediate-risk disease. This enrichment was unlikely to result from differences in EV recovery or technical variability, as EV-associated markers CD9 and CD63 were consistently detected across cohorts. Instead, the increased abundance of GD2-positive EVs likely reflects the biological characteristics of high-risk GD2-expressing neuroblastoma. This interpretation is supported by our *in vitro* findings demonstrating that EV-associated GD2 levels reflect GD2 expression on donor neuroblastoma cells. The simplicity of the bead-based workflow, which does not require ultracentrifugation or size-exclusion chromatography, further supports its potential suitability for future translational studies involving minimally invasive liquid biopsy samples.

The presence of GD2-positive EVs in patient plasma provides complementary information to established approaches for disease assessment. Their abundance showed only partial concordance with current biomarkers, suggesting that EV surface profiling captures a distinct biological dimension of neuroblastoma. Conventional serum biomarkers primarily reflect tumor burden or cellular turnover, whereas imaging provides anatomical information. In contrast, EV release is an active biological process influenced by viable tumor cells, membrane remodeling and interactions with the tumor microenvironment. The association between GD2-positive EV abundance and the mIBG soft tissue SIOPEN score may therefore reflect differences in tumor biology, vascular accessibility or EV release dynamics between disease compartments. Although these mechanisms remain to be defined, the incomplete overlap with established biomarkers supports the concept that EVs provide complementary rather than redundant information.

The complementary biological information provided by EV surface profiling is further illustrated by comparison with circulating cfDNA, which has emerged as a sensitive liquid biopsy approach in neuroblastoma. Circulating tumor DNA (ctDNA) enables detection of tumor-specific genomic alterations, intratumoral heterogeneity and molecular evolution during treatment. Studies by Schleiermacher and colleagues demonstrated that ctDNA can capture genomic heterogeneity and treatment-associated clonal evolution in neuroblastoma.^47,48^ We subsequently showed the utility of circulating cfDNA for longitudinal monitoring of clinically relevant genomic alterations.^11,49^ More recent studies demonstrated that serial cfDNA profiling can identify genetic and epigenetic features associated with tumor evolution and relapse^50^, while patient-specific *TERT* rearrangement breakpoints have been quantified longitudinally in ctDNA for monitoring treatment response and minimal residual disease.^10^ In addition, ctDNA sequencing has revealed emerging *ALK* alterations associated with treatment resistance in relapsed neuroblastoma.^51^ Multiplex biomarker approaches further highlight the value of combining complementary molecular measurements to resolve heterogeneous residual neuroblastoma cell populations within the bone marrow niche.^52^ In our pilot longitudinal analysis, ctDNA demonstrated a broader dynamic range and greater temporal variation than EV surface markers on the CD81^+^ EV population characterized, consistent with its established sensitivity for detecting genomic alterations. EVs, however, provide complementary biological information by reflecting the membrane phenotype of released vesicles, a feature not captured by DNA-based approaches. The detection of GD2-positive EVs in the earliest available plasma samples demonstrates the feasibility of identifying tumor-associated EV surface signatures at diagnosis. Although this pilot cohort was not designed to determine clinical sensitivity, prognostic value or treatment-response prediction, these observations support the concept that EV surface profiling and ctDNA interrogate different biological aspects of disease. Whereas ctDNA primarily reflects tumor genetics and clonal evolution, EV surface markers provide information on tumor-associated membrane characteristics. Integration of these approaches may therefore enable a more comprehensive assessment of disease status by combining genomic and phenotypic information.

Beyond their potential as biomarkers, GD2-positive EVs may contribute to tumor-immune interactions. Gangliosides influence membrane organization, receptor clustering and communication between tumor cells and immune components, suggesting that EV-associated GD2 may affect vesicle biodistribution, uptake or signaling in recipient cells. Tumor-derived EVs are increasingly recognized as regulators of immune responses and have been implicated in tumor-associated immune modulation.^53^ In addition, GD2 expression has been linked to immune escape mechanisms in neuroblastoma.^54,55^ The enrichment of GD2-positive EVs in high-risk neuroblastoma therefore raises the possibility that these vesicles contribute to systemic immune regulation. However, this remains a hypothesis requiring functional validation, including studies investigating the interactions between GD2-positive EVs and immune effector cells. The presence of GD2 on circulating EVs may also have implications for GD2-directed immunotherapies. Tumor-derived EVs have been shown to impair CAR T-cell activity against other tumor-associated antigens, including L1CAM^56^ and CD19^57^. Conversely, recent work demonstrated that GD2-positive EVs can activate GD2 CAR T-cells and enhance recognition of GD2-low neuroblastoma cells following EV exposure in vitro.^58^ The clinical consequences of circulating GD2-positive EVs during GD2-targeted therapy remain unknown. One possibility is that high levels of GD2-positive EVs may influence antibody-based therapies, including dinutuximab beta, by acting as circulating antigen reservoirs and thereby affecting antibody distribution and thereby affecting antibody distribution and antigen targeting on neuroblastoma cells. Alternatively, EV-associated GD2 may serve as a biomarker reflecting the presence of GD2-expressing disease and could support patient stratification for GD2-targeted approaches. Prospective studies integrating EV measurements with treatment response data will be required to determine whether EV-associated GD2 has predictive and/or pharmacodynamic relevance.

The analysis of EV surface markers remains technically challenging because EVs are heterogeneous, small and contain limited amounts of surface material. In this study, population-level EV surface profiling was mainly performed using the Miltenyi Biotec MACSPlex EV Kit Neuro, providing comparative information regarding surface-marker abundance across EV populations. However, this approach does not resolve individual-vesicle heterogeneity or determine marker co-expression patterns. The single EV analyses reported demonstrate GD2 detection on sEVs derived from IMR-5/75 and SK-N-BE(2) wildtype cells and a colocalization with the established tetraspanin EV markers CD63 or CD81. These findings support the potential value of GD2 as a neuroblastoma-associated EV surface marker. Flow cytometry-based methods on plasma EVs enable multiparametric analysis of individual EVs but require highly sensitive instrumentation, appropriate controls and standardized reporting due to limitations related to EV size and low signal intensity.^59,60^

Single-EV technologies, including high-sensitivity flow cytometry and interferometric approaches such as single-particle interferometric reflectance imaging sensing (SP-IRIS), have further advanced the characterization of EV subsets defined by specific surface-marker combinations.^60–62^ Standardized reporting frameworks, including MIFlowCyt-EV, will remain important for improving reproducibility and interpretation of EV measurements.^59,60^ Future studies integrating population-level profiling with single-EV approaches will be important to determine whether GD2-positive EVs represent a relatively homogeneous population or comprise distinct vesicle subsets with different biological properties.

Combining EV surface phenotyping with molecular cargo analysis, including RNA, DNA and protein profiling, may further improve specificity and enable identification of EV signatures associated with disease progression, treatment response or relapse. Such multimodal approaches may provide a more comprehensive representation of tumor biology than any individual biomarker class alone.

The present study establishes the presence and detectability of GD2-positive EVs in neuroblastoma but does not define their functional contribution to tumor progression, immune modulation or therapeutic response. Functional studies will be required to determine whether GD2-positive EVs directly influence tumor-immune interactions or treatment sensitivity. Furthermore, the longitudinal analysis was performed in a limited pilot cohort and should therefore be considered hypothesis-generating. Larger prospective studies with serial sampling during treatment and follow-up will be required to define the clinical utility of GD2-positive EVs for monitoring treatment response, minimal residual disease and relapse.

In conclusion, this study demonstrates that neuroblastoma-derived EVs preserve a characteristic tumor-associated surface phenotype comprising both membrane proteins and the clinically relevant glycolipid GD2. The identification of GD2-positive EVs in patient plasma highlights the EV surface glycolipidome as an underexplored layer of EV biology and provides a potential basis for expanding liquid biopsy strategies beyond conventional molecular analytes. Rather than replacing established biomarkers such as ctDNA, EV surface profiling offers complementary information by capturing membrane-associated features of tumor biology. Future integration of EV surface glycolipid and protein profiling with genomic biomarkers and advanced imaging approaches may provide a more comprehensive framework for precision monitoring and treatment stratification in neuroblastoma.

## Data availability

Original bead-based flow cytometry data reported in this study are available upon request from the corresponding author.

## Ethics approval

Biomaterial collection from patients with neuroblastoma was approved by EA2/055/17 and EA2/011/23. The surplus biomaterial usage from pediatric controls was permitted through EA2/131/11 and EA2/011/23.

## Patient consent

Written informed consent was obtained from all participants and/or their parents prior biomaterial storage and usage.

## Permission to reproduce material from other sources

Not applicable.

## Clinical trial registration

Patients were enrolled in the GPOH NB2016 Registry.

## ACKNOWLEDGEMENTS

The authors thank the patients and their parents, who agreed to take part in this study, and all physicians involved in biosample collection. The authors thank Jasmin Wünschel, Robin Gottschalk and Anika Winkler for technical support and the CECAD Imaging Facility in Cologne for transmission electron microscopy support. The authors thank Michael Mülleder and Li Jang (Department of Biochemistry, Charité – Universitätsmedizin Berlin), Anja Heeren-Hagemann and Nicole Huebener (Department of Pediatric Oncology and Hematology, Charité – Universitätsmedizin Berlin) as well as Carsten Geiß (Department of Anesthesiology, University Medical Center of Johannes Gutenberg University Mainz) for helpful discussions and critical feedback on the manuscript.

