## supplementary figures S1 - S4 for "Circulating extracellular vesicles display immunotherapeutic targets and complement circulating tumor DNA in high-risk neuroblastoma"

**Table of Contents**

**Supplementary Figures** starting p. 2

**Legends to Supplementary Figures** starting p. 8

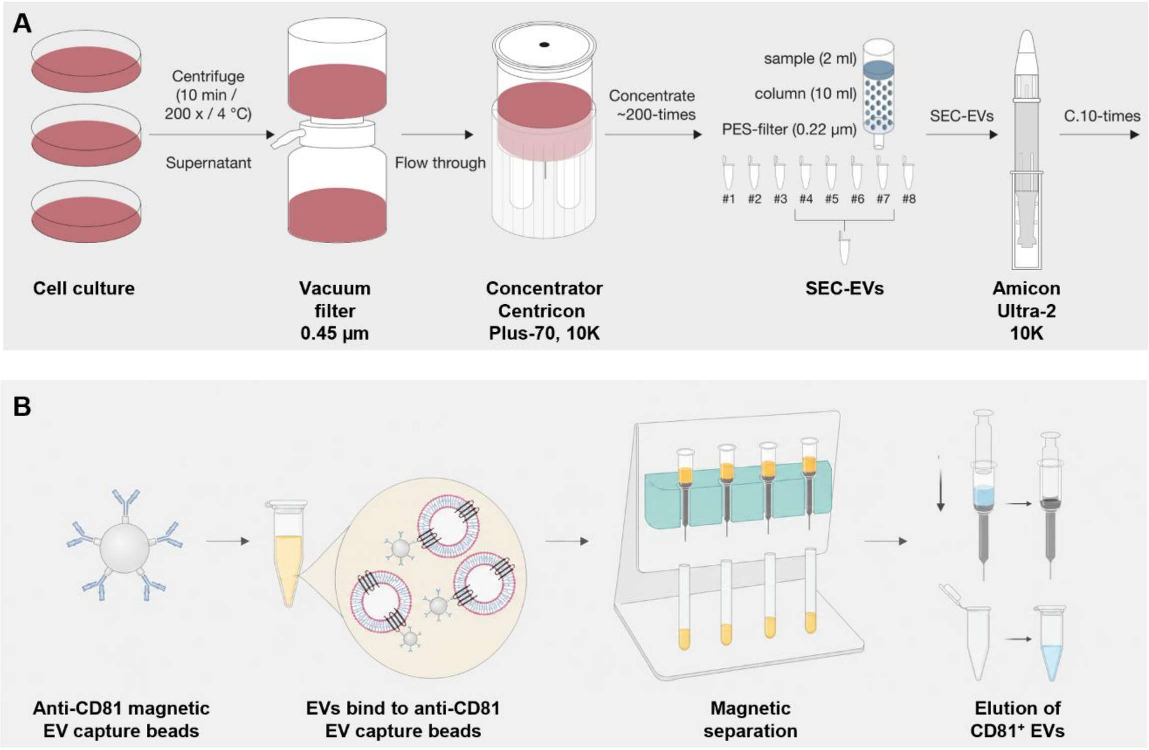


**Suppl. Fig. S1. Workflow for extracellular vesicle (EV) isolation from cell culture medium and human blood plasma. A**, Small EVs isolated from conditioned cell culture medium by size-exclusion chromatography (SEC) for *in vitro* experiments. **B**, EVs isolated from human blood plasma by immunocapture using magnetic beads conjugated to an antibody against CD81 for clinical analyses.

**A - IMR-5/75 sEVs**

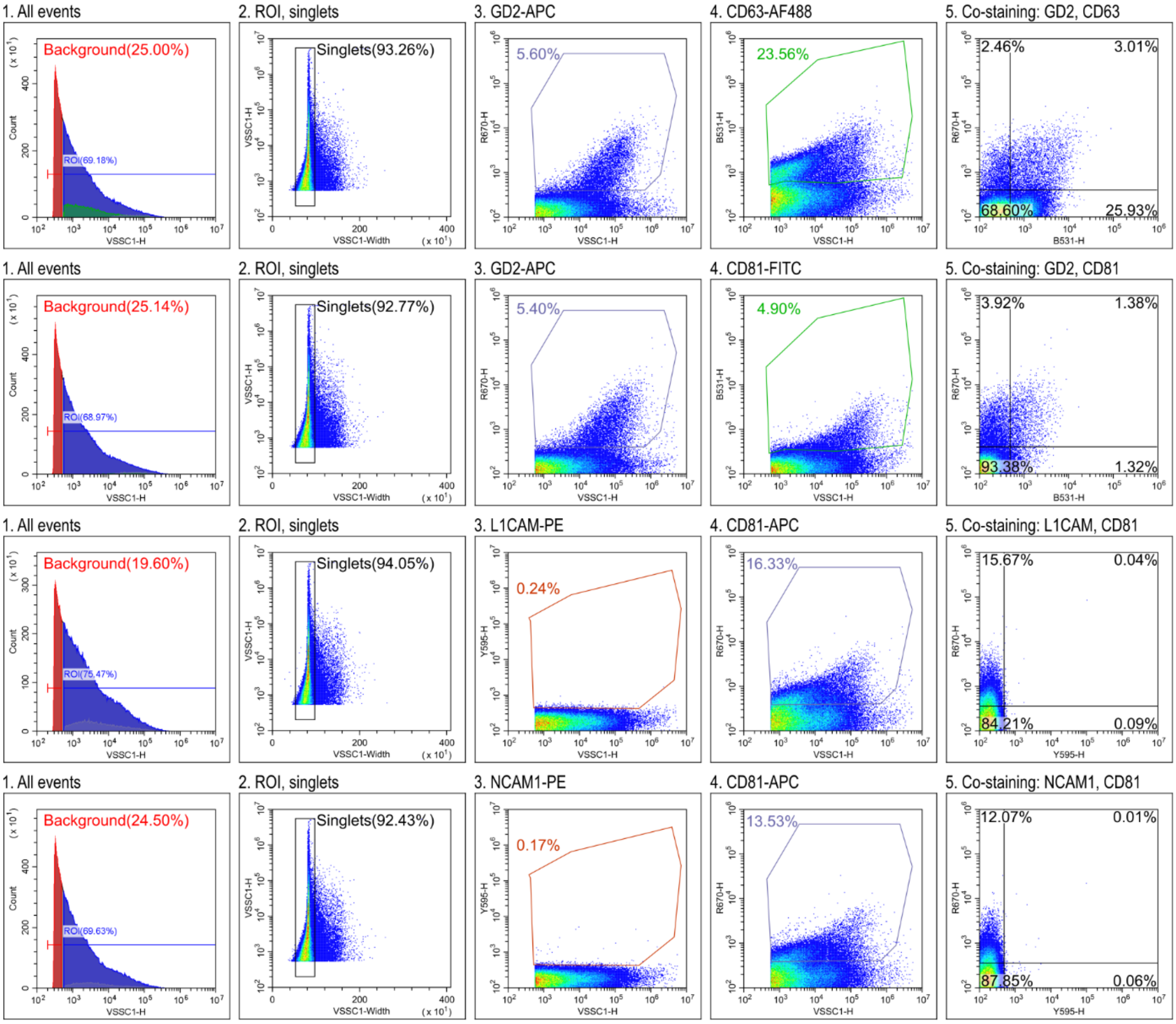


**B - SK-N-BE(2) wildtype sEVs**

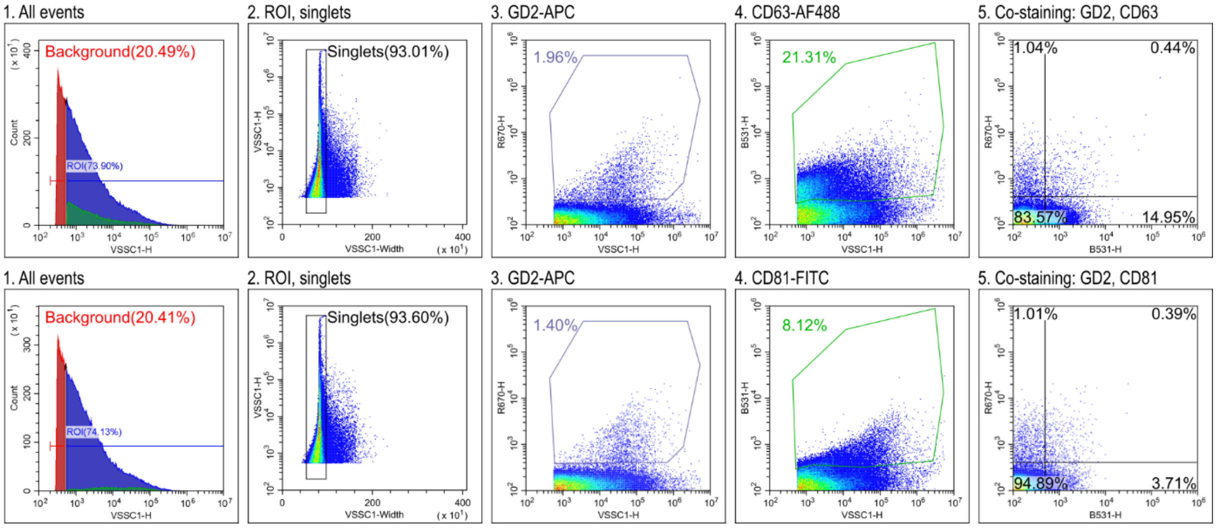

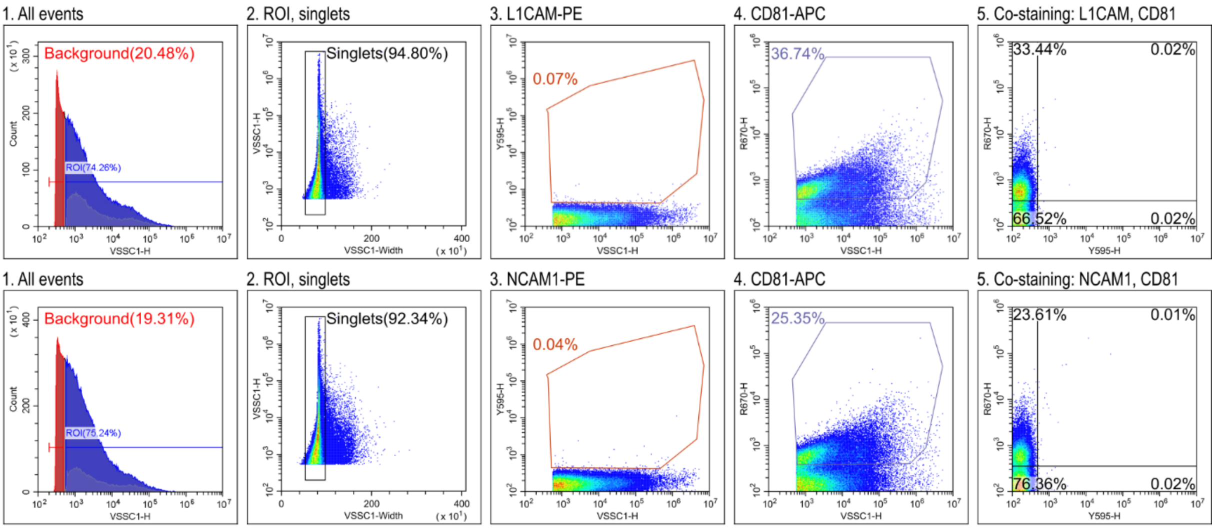


**C - SK-N-BE(2) *B4GALNT1*[knockout] sEVs**

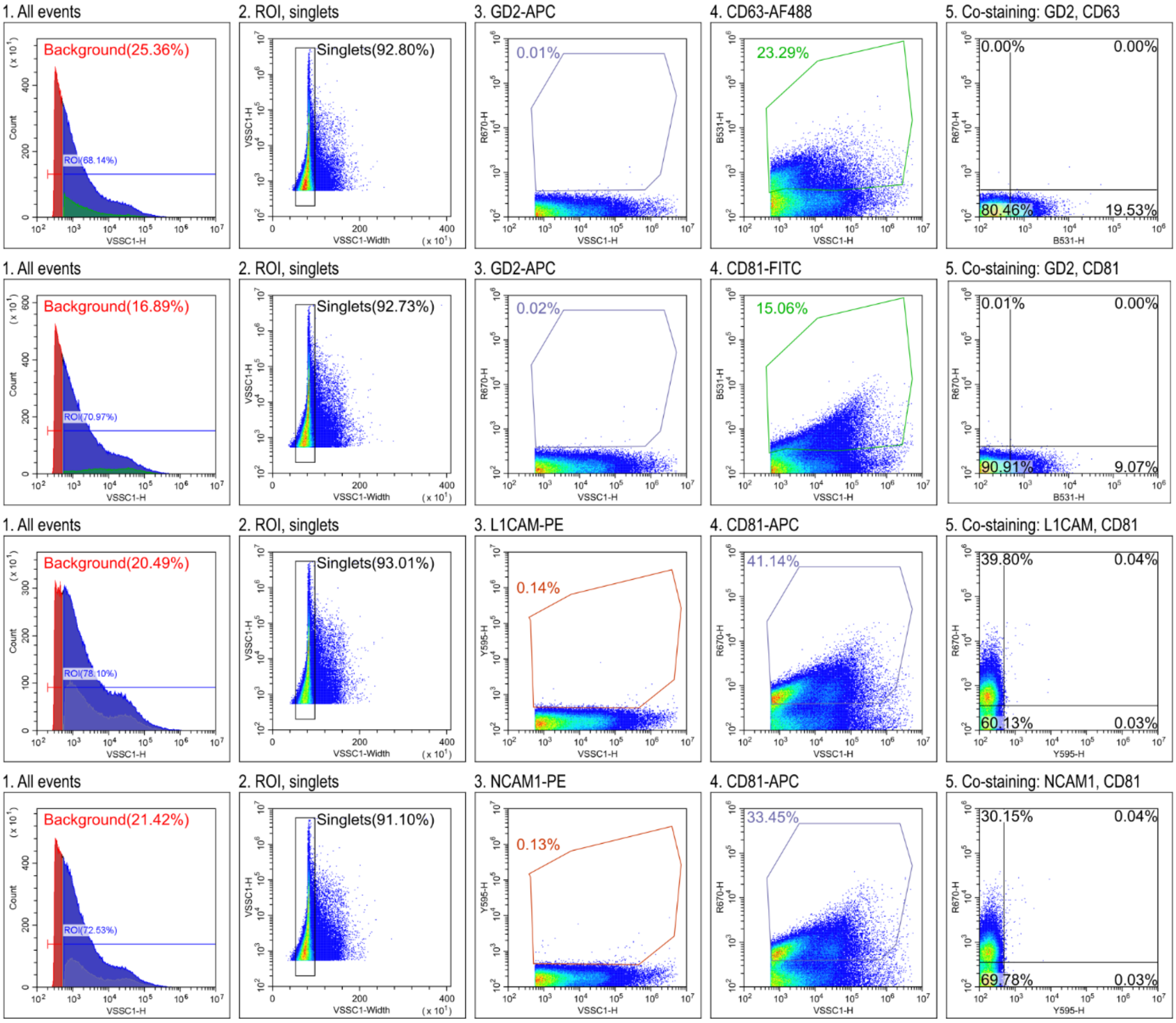


**Suppl. Fig. S2. Gating strategy for single vesicle flow cytometry analysis.** Small extracellular vesicles (sEVs) from the human neuroblastoma cell lines IMR-5/75 (**A**), SK-N-BE(2) (**B**) and the GD2-deficient SK-N-BE(2) *B4GALNT1* knockout model (**C**) were isolated from conditioned cell culture medium by size-exclusion chromatography and stained with the indicated antibodies. The gating strategy is illustrated in the numbered panels: background signals were first excluded (1), followed by singlet selection within the region of interest (ROI, 2). Marker-positive gates were defined using unstained controls, and marker-positive events are shown in panels

3 and 4. Events positive for both markers in the respective co-staining analyses are shown in panel 5. The detectors correspond to the following fluorophores: Y595-H detects PE, R670-H detects APC and B531-H detects AF488 and FITC.

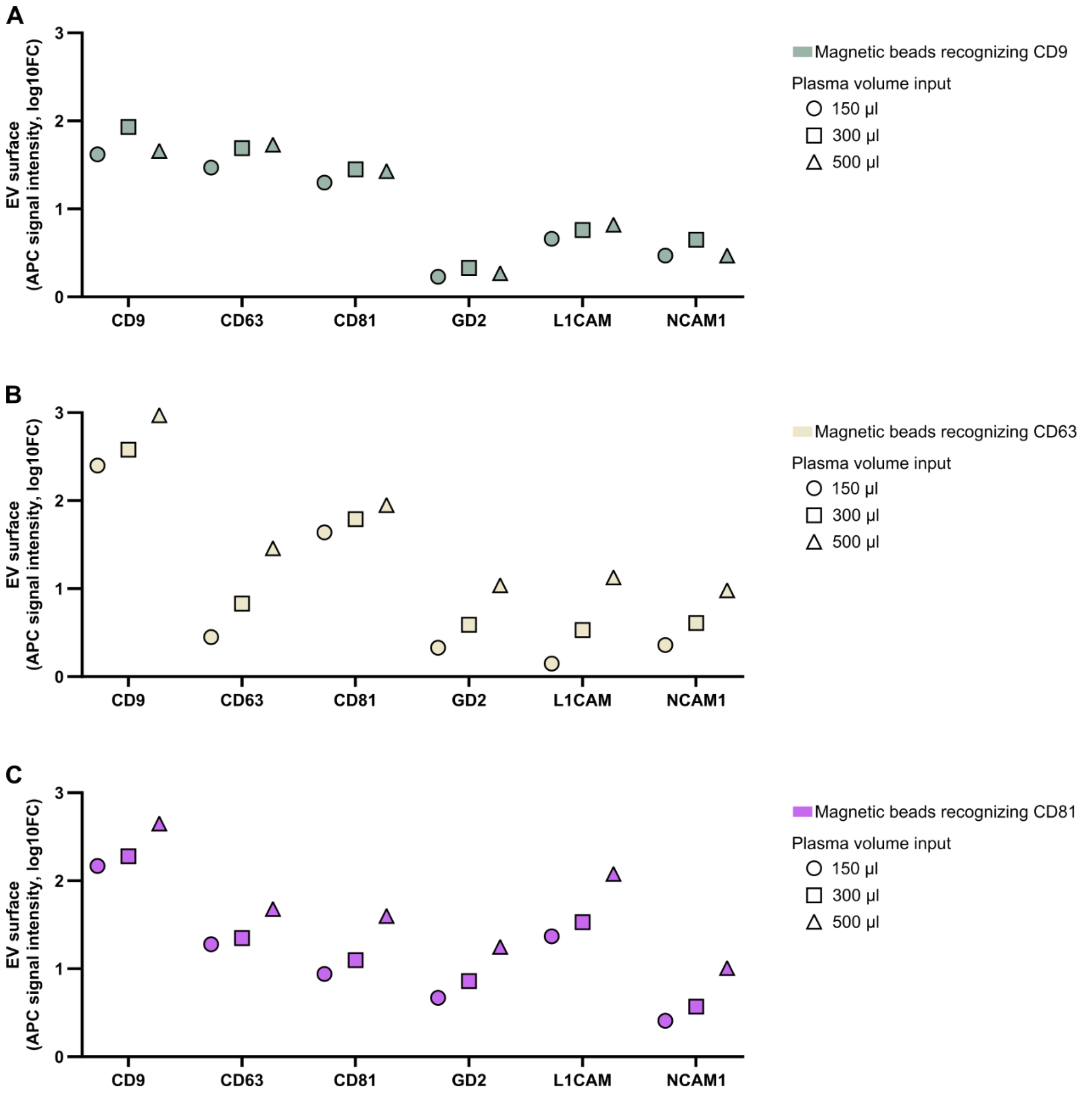


**Suppl. Fig. S3. Comparison of magnetic bead-based isolation of EVs from blood plasma samples**. Magnetic beads coated with anti-CD9 (**A**), anti-CD63 (**B**) or anti-CD81 (**C**) were used to purify EVs from a blood plasma sample from a patient with high-risk neuroblastoma. The tetraspanins used for EV capture show reduced signal intensities, because the corresponding epitopes are mainly masked by the antibodies used for bead-based isolation. In total, three different plasma volumes were assessed. Analyses employing the Miltenyi Biotec MACSPlex EV Kit Neuro included the respective tetraspanins as well as the neuroblastoma associated markers GD2, L1CAM and NCAM1. Data are shown as the log10-fold change in allophycocyanin (APC) signal intensity relative to buffer control.

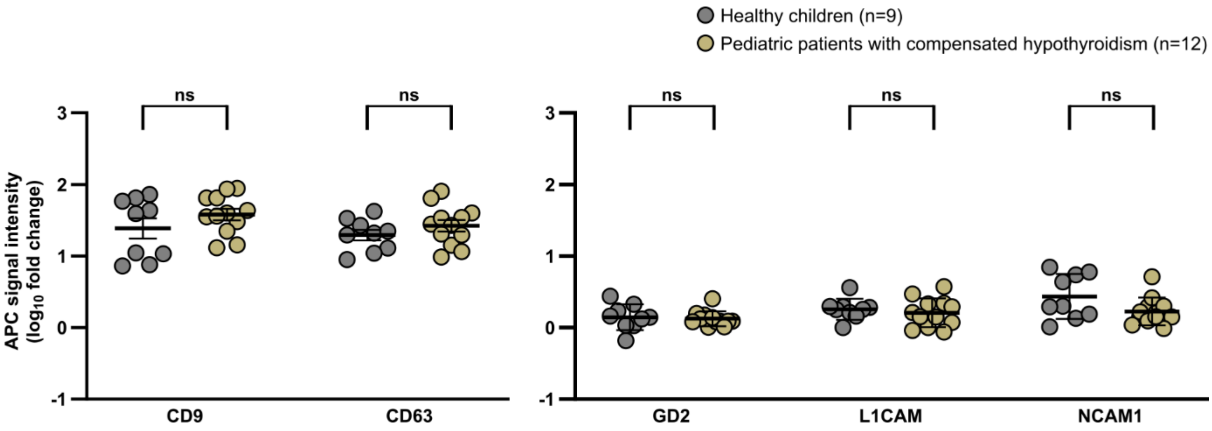


**Suppl. Fig. S4. Flow-cytometric analysis of CD9, CD63, GD2, L1CAM and NCAM1-positive EVs in human blood plasma from pediatric controls.** CD9 and CD63 expression on EVs is shown in the *left panel*, whereas GD2, L1CAM and NCAM1-positive EVs are shown in the *right panel*. EVs were isolated from blood plasma using CD81-targeting magnetic beads and analyzed by flow cytometry. The cohort comprised healthy children (n = 9) and children with compensated hypothyroidism (n = 12). Data are shown as log10-fold change in allophycocyanin (APC) signal intensity relative to baseline, with individual measurements and arithmetic mean ± S.E.M. Statistical significance was assessed between the two groups. ns, not significant.

### LEGENDS TO SUPPLEMENTARY FIGURES

**Suppl. Fig. S1. Workflow for extracellular vesicle (EV) isolation from cell culture medium and human blood plasma. A**, Small EVs isolated from conditioned cell culture medium by size-exclusion chromatography (SEC) for *in vitro* experiments. **B**, EVs isolated from human blood plasma by immunocapture using magnetic beads conjugated to an antibody against CD81 for clinical analyses.

**Suppl. Fig. S2. Gating strategy for single vesicle flow cytometry analysis.** Small extracellular vesicles (sEVs) from the human neuroblastoma cell lines IMR-5/75 (**A**), SK-N-BE(2) (**B**) and the GD2-deficient SK-N-BE(2) B4GALNT1 knockout model (**C**) were isolated from conditioned cell culture medium by size-exclusion chromatography and stained with the indicated antibodies. The gating strategy is illustrated in the numbered panels: Background signals were first excluded (1), followed by singlet selection within the region of interest (ROI, 2). Marker-positive gates were defined using unstained controls, and marker-positive events are shown in panels 3 and 4. Events positive for both markers in the respective co-staining analyses are shown in panel 5. The detectors correspond to the following fluorophores: Y595-H detects PE, R670-H detects APC and B531-H detects AF488 and FITC.

**Suppl. Fig. S3. Comparison of magnetic bead-based isolation of EVs from blood plasma samples.** Magnetic beads coated with anti-CD9 (**A**), anti-CD63 (**B**) or anti-CD81 (**C**) were used to purify EVs from a blood plasma sample from a patient with high-risk neuroblastoma. The tetraspanins used for EV capture show reduced signal intensities, because the corresponding epitopes are mainly masked by the antibodies used for bead-based isolation. In total, three different plasma volumes were assessed. Analyses employing the Miltenyi Biotec MACSPlex EV Kit Neuro included the respective tetraspanins as well as the neuroblastoma associated markers GD2, L1CAM and NCAM1. Data are shown as the log10-fold change in allophycocyanin (APC) signal intensity relative to buffer control.

**Suppl. Fig. S4. Flow-cytometric analysis of CD9, CD63, GD2, L1CAM and NCAM1-positive EVs in human blood plasma from pediatric controls.** CD9 and CD63 expression on EVs is shown in the left panel, whereas GD2, L1CAM and NCAM1-positive EVs are shown in the right panel. EVs were isolated from blood plasma using CD81-targeting magnetic beads and analyzed by flow cytometry. The cohort comprised healthy children (n = 9) and children with compensated hypothyroidism (n=12). Data are shown as log10-fold change in allophycocyanin (APC) signal intensity relative to baseline, with individual measurements and arithmetic mean ± S.EM. Statistical significance was assessed between the two groups. ns, not significant.
